# Isolation and structure of the fibril protein, a major component of the internal ribbon for *Spiroplasma* swimming

**DOI:** 10.1101/2021.02.24.432793

**Authors:** Yuya Sasajima, Takayuki Kato, Tomoko Miyata, Akihiro Kawamoto, Keiichi Namba, Makoto Miyata

**Affiliations:** Graduate School of Science, Osaka Metropolitan University, Osaka, Japan; Institute for Protein Research, Osaka University, 3-2 Yamadaoka, Suita, Osaka, Japan; Graduate School of Frontier Biosciences, Osaka University, 1-3 Yamadaoka, Suita, Osaka, Japan; RIKEN Center for Biosystems Dynamics Research and Spring-8 Center, 1-3 Yamadaoka, Suita, Osaka, Japan; JEOL YOKOGUSHI Research Alliance Laboratories, Osaka University, 1-3 Yamadaoka, Suita, Osaka, Japan; The OCU Advanced Research Institute for Natural Science and Technology (OCARINA), Osaka Metropolitan University, Osaka, Japan

**Author notes:** **Correspondence:** Makoto Miyata.

**Keywords:** Helical shape, Motility, Cytoskeleton, Filament, Single particle analysis, Quick freeze replica electron microscopy, Electron tomography

## Abstract

*Spiroplasma*, which are known pathogens and commensals of arthropods and plants, are helical-shaped bacteria that lack a peptidoglycan layer. *Spiroplasma* swim by alternating between left- and right-handed helicity. Of note, this system is not related to flagellar motility, which is widespread in bacteria. A helical ribbon running along the inner side of the helical cell should be responsible for cell helicity and comprises the bacterial actin homolog, MreB, and a protein specific to *Spiroplasma*, fibril. Here, we isolated the ribbon and its major component, fibril filament, for electron microscopy (EM) analysis. Single-particle analysis of the fibril filaments using the negative-staining EM revealed a three-dimensional chain structure composed of rings with a size of 11 nm wide and 6 nm long, connected by a backbone cylinder with an 8.7 nm interval with a twist along the filament axis. This structure was verified through EM tomography of quick-freeze deep-etch replica sample, with a focus on its handedness. The handedness and pitch of the helix for the isolated ribbon and fibril filament agreed with those of the cell in the resting state. Structures corresponding to the alternative state were not identified. These results suggest that the helical cell structure is supported by fibril filaments; however, the helical switch is caused by the force generated by the MreB proteins. The isolation and structural outline of the fibril filaments provide crucial information for an in-depth clarification of the unique swimming mechanism of *Spiroplasma*.

## 1 Introduction

Mollicutes, which are parasitic or commensal bacteria, evolved from the phylum, Firmicutes, including *Bacillus* and *Clostridium* by reducing their genome size (Razin et al., 1998;Razin and Hayflick, 2010;Grosjean et al., 2014;Miyata et al., 2020). During the course of evolution, the cells became softer and smaller owing to the loss of the peptidoglycan layer. These changes may have allowed some species to transmit their internal housekeeping activities, such as the rotation of ATP synthesis to the outside, resulting in the acquisition of at least three unique motility mechanisms (Relich et al., 2009;Miyata and Hamaguchi, 2016a;b;Distelhorst et al., 2017;Miyata et al., 2020;Toyonaga et al., 2021). Two of the three well studied mechanisms are exerted by *Mycoplasma mobile* and *Mycoplasma pneumoniae*. These species exhibit gliding motilities on solid surfaces, in which leg structures repeatedly catch sialylated oligosaccharides on host cells based on two mechanisms (Miyata, 2010;Miyata and Hamaguchi, 2016a;b). Another motility system is the helicity-switching swimming of *Spiroplasma*, which is the subject of the present study (Movie_S1) (Shaevitz et al., 2005;Wada and Netz, 2009;Harne et al., 2020b;Sasajima and Miyata, 2021). *Spiroplasma* species are parasitic to plants and arthropods and are characterized as polarized helical-shaped cells with one tapered end (Gasparich, 2002;Harumoto and Lemaitre, 2018;Harne et al., 2020b). These species exhibit obvious chemotaxis despite the absence of genes for the two-component regulatory system in the genome, which is generally responsible for bacterial chemotaxis (Liu et al., 2017). In general, swimming bacteria, including spirochetes, can migrate through the rotational motion of the flagellar motor fixed to the peptidoglycan layer, whereas *Spiroplasma* has a unique swimming system in which kinks propagate along the cell body with a switch between left- and right-handed cell helicity (Fig. 1A). The outline of this mechanism has been clarified as follows. The rotation of helical cells linked to the helicity switch pushes the water back (Trachtenberg and Gilad, 2001;Trachtenberg et al., 2003b;Kürner et al., 2005;Shaevitz et al., 2005;Wada and Netz, 2009;Sasajima and Miyata, 2021). The helicity might be dominated by an intracellular structure called the “ribbon,” which localizes along the innermost line of the helical cell structure and is composed of protofilaments. Based on structural studies, ribbons may switch their helicity through changes in the protofilament length (Trachtenberg and Gilad, 2001;Kürner et al., 2005;Cohen-Krausz et al., 2011). Ribbons are known to be composed of fibril proteins specific for *Spiroplasma* species and some *Spiroplasma* MreB (SMreB) proteins related to MreB that are common in rod-shaped bacteria. Although fibril filaments are featured by repetitive ring structures, nanometer-order three-dimensional structure has not been clarified (Trachtenberg and Gilad, 2001;Trachtenberg et al., 2003a;Kürner et al., 2005;Trachtenberg et al., 2008;Cohen-Krausz et al., 2011;Liu et al., 2017).

**Figure 1.**
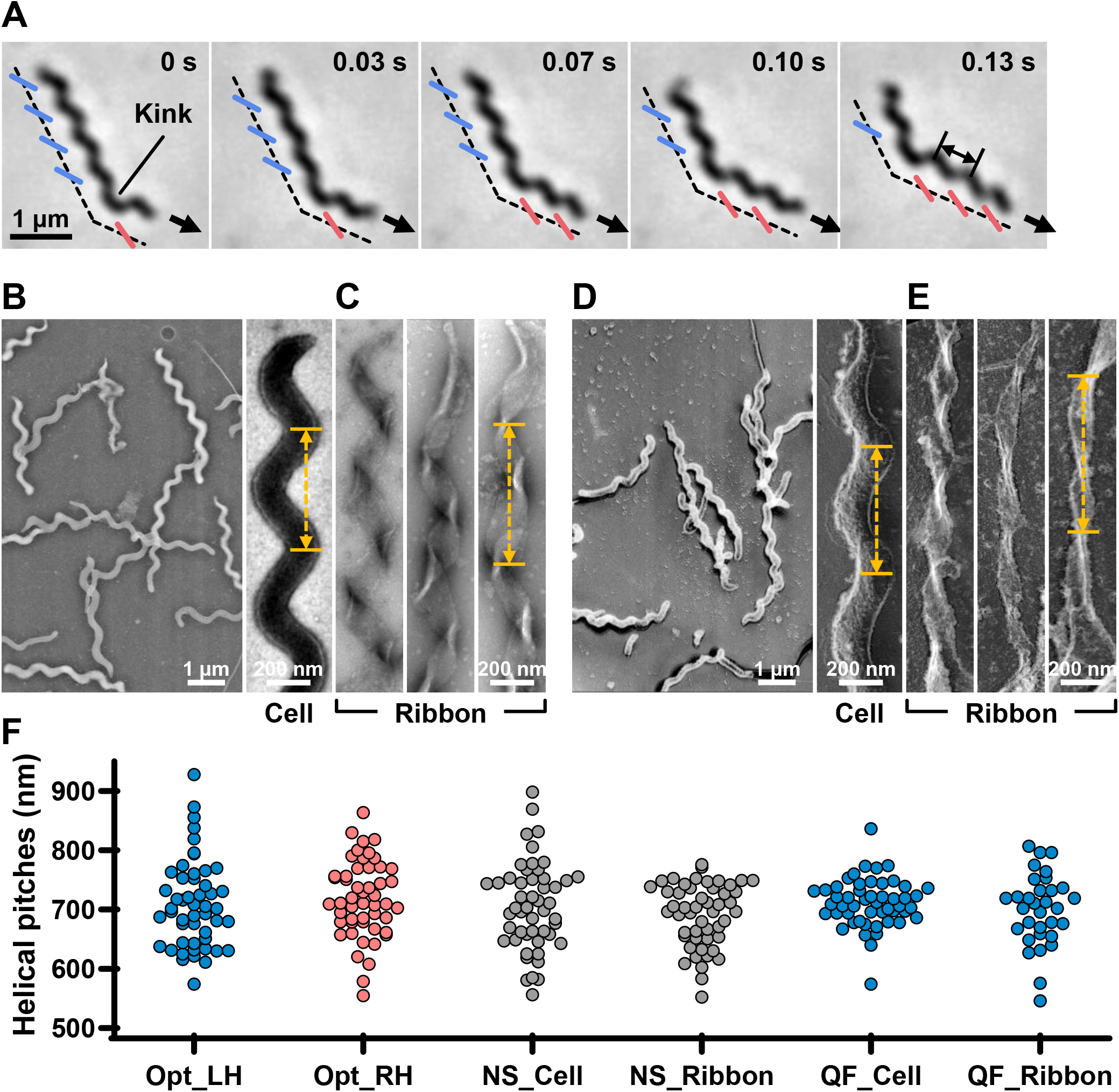
Helicity of the cell and ribbon structure. **(A)** Phase-contrast microscopy of swimming cell. The blue and red segments, and broken line indicate the left- and right-handed helicity, and cell axes, respectively. The pitch was measured as indicated by a double headed arrow. **(B, C)** Cell and ribbon images acquired by negative-staining EM. **(D, E)** Cell and ribbon images acquired by QFDE EM. (F) Helical pitches of cells and ribbon measured by optical microscopy, negative-staining EM, and QFDE-EM. Handedness was judged by optical microscopy and QFDE-EM. All cells analyzed by QFDE-EM were left-handed as they were grown under a starved condition.

In the present study, we isolated the filament of fibrils, the major component protein of ribbons and clarified its nanometer-order three-dimensional structure using electron microscopy (EM) and image analyses. The fibril filament has a repetitive structure featuring a ring and a cylinder with a helical pitch similar to those of the ribbon and cell.

## 2 Materials and Methods

### 2.1 Bacterial strains and culture conditions

The type strain, TDA-040725-5^T^, of *Spiroplasma eriocheiris* was cultured in R2 medium (2.5% [wt/vol] heart infusion broth, 8% sucrose, and 10% horse serum) at 30 °C until an optical density of 0.06 to 0.1 was achieved at 600 nm (Liu et al., 2017;Terahara et al., 2017).

### 2.2 Optical microscopy

Cultured cells were centrifuged at 11 000 × *g*, 10 °C for 10 min and suspended in PBS consisting of 75 mM sodium phosphate [pH 7.3], 100 mM NaCl containing 20 mM glucose, and 0.6% methylcellulose, to achieve a cell density 10-fold higher than that of the original (Liu et al., 2017;Terahara et al., 2017). Cells were inserted into a tunnel chamber assembled by taping coverslips, as previously described, and observed under an IX71 microscope (Olympus, Tokyo, Japan) (Uenoyama et al., 2004). A video was captured using a DMK33UX174 complementary metal–oxide–semiconductor (CMOS) camera (The Imaging Source, Taipei, Taiwan) and analyzed using ImageJ v1.53a (https://imagej.nih.gov/ij/).

### 2.3 Electron microscopy

To observe the intact cells, the cell suspension was placed on a hydrophilized grid, fixed using 2% glutaraldehyde, washed with water, and stained with 2% uranyl acetate. To observe the internal structure, the cell suspension on a grid was treated with PBS containing 0.1 mg/mL DNase and 1 mM MgCl_2_ for 20 s, washed, and stained with 2% uranyl acetate. QFDE-EM was performed as previously reported for specimens suspended in a solution, 10 mM HEPES (pH 7.6), and 150 mM NaCl containing mica flakes (Tulum et al., 2019). The Triton X-100 treatment was done on glass surface before freezing, to observe the internal structure. Images were acquired using a JEM1010 EM (JEOL, Akishima, Japan) equipped with a FastScan-F214(T) charge-coupled device (CCD) camera (TVIPS, Gauting, Germany) and analyzed using ImageJ v1.53a. For tomography, images were captured using a Talos F200C EM (FEI, Eindhoven, Netherlands) equipped with a 4k × 4 K Ceta CMOS camera (FEI). Single-axis tilt series were collected covering an angular range from -50° to +50° with 1.5° steps and analyzed using IMOD (ver 4.11) and PEET (ver 1.15.0).

### 2.4 Isolation of the ribbon and fibril

To isolate the internal structure, 10 mL of cell suspension in PBS was treated with 1% Triton X-100, 0.1 mg/mL DNase, 1 mM MgCl_2_, and 0.1 mM PMSF, with shaking for 10 min at 4 °C. The insoluble fraction was recovered via centrifugation at 20 000 × *g* for 30 min at 4 °C, and suspended in PBS to obtain a final volume of 0.2 mL. The sample was placed at the top of sucrose solution layers of 0%, 20%, 30%, 40%, 50%, and 60%, and centrifuged at 20 000 × *g* for 20 min at 4 °C in a 1.5 mL tube at a fixed angle. To isolate the fibril filament, the insoluble fraction was additionally treated with a solution consisting of 2% cholic acid, 20 mM Tris-Cl pH 8.0, 150 mM NaCl at 4 °C for 8 h, and subjected to stepwise density gradient centrifugation. SDS-PAGE and peptide mass fingerprinting were performed as described previously (Nakane and Miyata, 2007;Kawakita et al., 2016;Liu et al., 2017). Band intensities were calculated using ImageJ, from scanned gel images.

### 2.5 Preparation of the single-stranded fibril filament

The isolated fibril was adjusted to 1 mg/mL in 20 mM Tris-Cl pH 8.0 and 150 mM NaCl. The fibril suspension (1 mL in a 1.5 mL test tube) was treated on ice for 5 s using a sonicator (UR-21P, TOMY, Tokyo, Japan). The condition of the fibril filament was checked via negative-staining electron microscopy (EM). The processes of sonication and observation were repeated with the fibril suspension until more than 90% of the filaments became single-stranded.

### 2.6 Reconstitution of the 3D structure

The contrast transfer function (CTF) parameters for negative-staining EM images were estimated using the Gctf25 software (Zhang, 2016). The images of fibril filaments were selected automatically using RELION 3.0 (Zivanov et al., 2018) as helical objects and segmented as squares of 200 × 200 pixels with a 90% overlap. These 14,543 images were 2D-classified and 11,867 images were selected for further analyses. *Ab-initio* reconstitution was performed using cisTEM (Grant et al., 2018) based on segmented images from 12 classes. The selected 11,867 particle images were 3D-classified using the 3D map in RELION 3.0 (Zivanov et al., 2018).

## 3 Results

### 3.1 Cell helicity is derived from the internal ribbon structure

To clarify the relationship between the helical cell morphology and the internal ribbon structure, we first measured the helical pitches of the swimming cells using optical microscopy. Under phase-contrast microscopy, the helical shape of the cells can be observed as a series of dense segments in the defocused image plane relative to the cell axis (Fig. 1A). We measured the pitches along the cell axis for the segments of left and right handedness (Fig. 1F). The helical pitches were 709 ± 74 (n = 50) and 718 ± 65 nm (n = 50) for the left- and right-handed segments, respectively.

EM was subsequently employed to analyze the internal ribbon structure and compare the helical pitches of the cells and ribbons. Negative-staining EM revealed images of helical-shaped cells with a narrow tip on one side (Fig. 1B).

The internal ribbon structure was exposed by treating the cells with 0.1% Triton X-100 on the grid (Fig. 1C). The ribbon had a “helical” flat structure. These observations are consistent with those of previous studies (Trachtenberg and Gilad, 2001). Thereafter, the pitches of the cell and the exposed ribbon were measured (Fig. 1F). Generally, the specimens for negative-staining EM are placed in vacuum and dried, which can result in distortions and is disadvantageous for evaluating the helicity. We therefore applied quick-freeze, deep-etch (QFDE) EM to visualize the structure in a state as closely as possible to the original (Heuser, 2011). In QFDE, a sample is frozen in milliseconds and exposed by fracturing and etching. Thereafter, a platinum replica was created by shadowing. The observation of the replica by transmission EM provides images with high contrast and resolution, which is markedly better than that provided by conventional scanning electron microscopy (SEM) (Heuser, 2011;Tulum et al., 2019). Replicas were then prepared by fracturing and platinum coating. QFDE-EM revealed cell morphology consistent with that obtained using negative-staining EM (Fig. 1D). Using QFDE-EM, we observed the ribbons exposed to 0.1% Triton X-100 treatment (Fig. 1E). The ribbon had a structure in which the twisted positions were assembled in a line, showing that the images observed by negative-staining EM were flattened. Interestingly, all ribbons were left-handed (Fig. 1D, E). When the cells were starved in phosphate-buffered saline (PBS) without glucose for 30 min, they showed a left-handed helix with the same pitch. Therefore, this structure was assumed to be the default state of the cell, and the ribbon switched to the default structure during the visualization process. The helical pitches of the cells and ribbons aligned well with each other, indicating that the ribbon has a critical role for cell helicity (Fig. 1F, Table 1).

**Table 1.** Dimensions of the cell and ribbon

| Parameters | Negative-staining<br>EM | QFDE-EM | Optical<br>microscopy |
| --- | --- | --- | --- |
| Cell helical pitch | 706 ± 74 nm<br>703 nm | 711 ± 41 nm (LH)<br>711 nm (LH) | 709 ± 74 nm (LH) |
|  |  |  | 702 nm (LH) |
|  |  |  | 718 ± 65 nm (RH) |
|  |  |  | 711 nm (RH) |
| Ribbon helical pitch | 691 ± 53 nm<br>700 nm | 700 ± 60 nm (LH)<br>706 nm (LH) |  |
| Isolated ribbon 1/2 helical<br>pitch | 350 ± 17 nm<br>352 nm |  |  |
| Isolated fibril 1/2 helical pitch | 341 ± 27 nm<br>335 nm | 351 ± 34 nm (LH)<br>352 nm (LH) |  |

Handedness is represented by LH and RH. The upper and lower rows show the mean, standard deviation, and median, respectively.

### 3.2 Isolation and characterization of the ribbon

For further characterization, we isolated the internal ribbon structure. The cell suspension was treated with 1% Triton X-100 and subjected to stepwise gradient centrifugation with 0%, 20%, 30%, 40%, 50%, and 60% sucrose layers. After centrifugation, we found a dense layer of cell contents at the bottom of the 40% sucrose layer. The fraction was recovered and then observed by EM. Based on the observation, the ribbon was found to comprise protofilaments with a width of 66 ± 12 nm (n=20) and a length longer than 2 μm, which may correspond to the full length of the cell (Fig. 2A). To analyze the number and width of the protofilaments involved in the isolated ribbon, we traced a cross sectional image profile of the ribbon (Fig. 2D a). Six to nine protofilaments were detected, with peak distances ranging between 4 and 16 nm (Fig. 2D b, c), consistent with the findings of the previous studies (Trachtenberg and Gilad, 2001;Liu et al., 2017). Ribbon twists are observed as periodic frays in the ribbons. The ribbon pitches were measured from the frays as 350 ± 17 nm (n = 47) (Fig. 2D d), which is comparable to the helical pitches of the cells and the ribbons exposed from cells on grids (Fig. 1, Table 1) (*P* = 0.7 > 0.01). SDS-PAGE and peptide mass fingerprinting analyses of this fraction revealed five protein bands, including six proteins (Fig. 2B, Table 2, Table S1). Band (v) contains SMreBs 2 and 4. The whole ribbon fraction mainly comprised the fibril protein (band iii) and a protein mixture of SMreBs 2 and 4 (band v), with an intensity ratio of 47% and 37% of the total protein amount, respectively. Further studies are necessary to conclude physical interactions of SPE-1201 and FtsH to fibril protein, because these proteins are abundant in *S eriocheiris* cells (Liu et al., 2017).

**Table 2.** Protein components of the ribbon isolated from original cells^1^.

| Protein<br>Band<sup>1</sup> | Gene ID | Annotation | Mascot<br>Score<sup>2</sup> | Mass<br>(kDa)<sup>3</sup> | Density ratio (%) |  |
| --- | --- | --- | --- | --- | --- | --- |
|  |  |  |  |  | Original | A22 treated |
| (i) | SPE-1201 | Hypothetical<br>protein | 72 | 85.8 | 4 | 5 |
| (ii) | SPE-0013 | FtsH | 84 | 77.0 | 12 | 17 |
| (iii) | SPE-0666 | Fibril | 206 | 58.7 | 47 | 67 |
| (iv) | SPE-1231 | SMreB5 | 98 | 38.5 | 10 | 7 |
| (v) | SPE-1224 | SMreB2 | 80 | 37.8 | 27 | 4 |
|  | SPE-1230 | SMreB4 |  | 40.7 |  |  |

<sup>1</sup> From A22-treated cells, the proteins common to the original cells were identified for bands (i)–(iv). For band (v) only, SMreB2 was identified.
<sup>2</sup> Mascot Score is the logarithm of probability that the observed match is a random event.
<sup>3</sup> Calculated from the amino acid sequence as a monoisotopic molecule.

**Figure 2.**
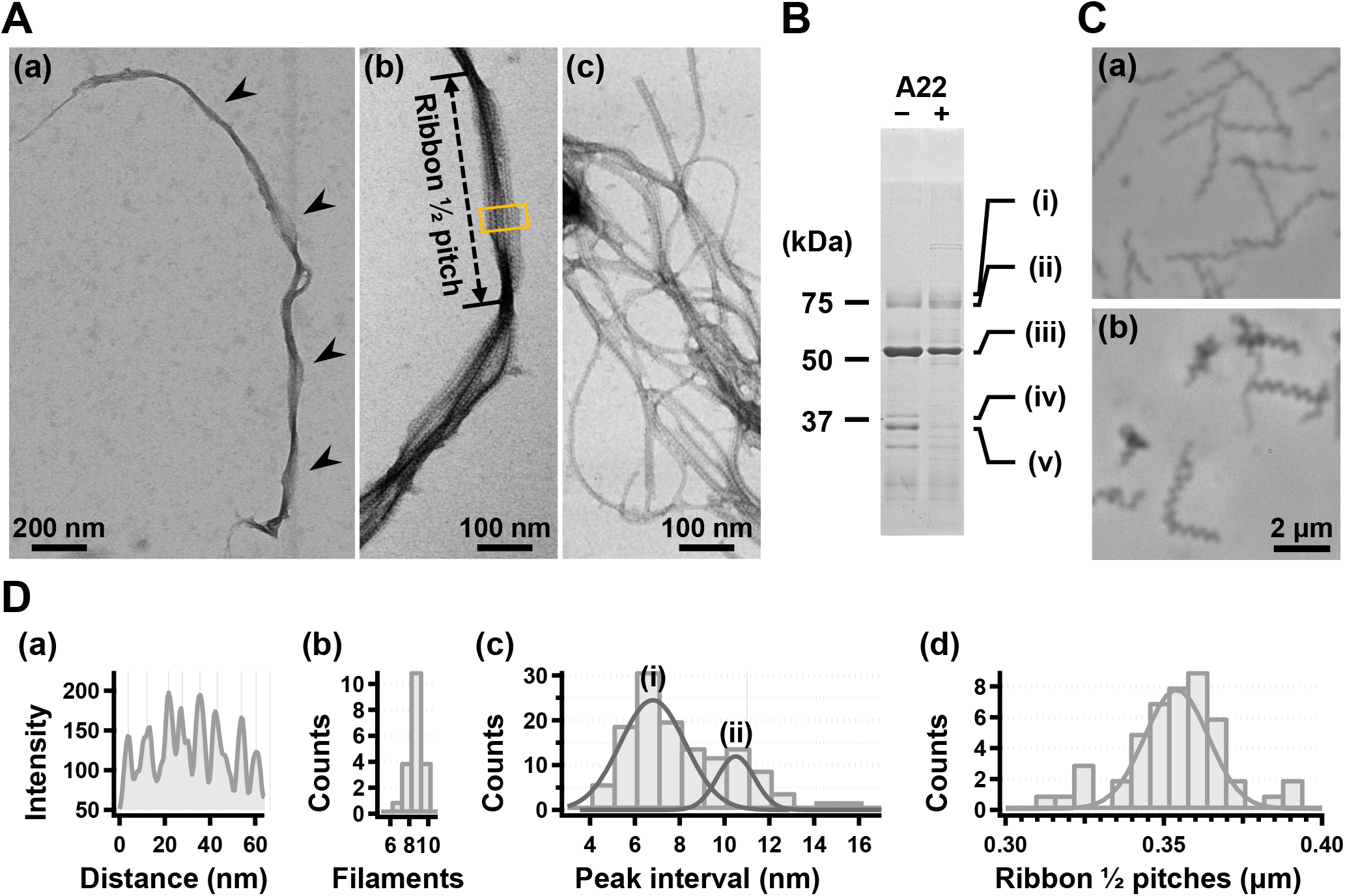
Isolation and characterization of the ribbon. **(A)** Isolated ribbon structure. (a) The whole structure of the isolated ribbon with helicity as shown by periodical wide positions (marked by arrows). (b) The magnified image of the isolated ribbon and the helical pitch is indicated by a bidirectional arrow. (c) Ribbon fraction isolated from cells treated with A22. **(B)** Protein profiles of the ribbon fraction isolated from cells untreated and treated with A22. **(C)** Cell images before (a) and after (b) treatment with 1 mM A22 for 2 min. **(D)** Numerical characterization. (a) Sectional image profile of the area boxed in panel (A b). The peaks correspond to the center of the protofilament. (b) Histogram for the number of protofilaments involved in a ribbon. (c) Histogram for the protofilament width in ribbons. The distribution can be fitted by two Gaussian curves marked (i) and (ii), with peaks around 7.0 and 10.5 nm, respectively. (d) Histogram for the helical pitches of the isolated ribbon, fitted by a Gaussian curve with a peak at 351 ± 16 nm (n=47).

We intended to use A22, an inhibitor of MreB polymerization, to examine the role of SMreBs in ribbon formation (Shi et al., 2018); this is because the binding of A22 to SMreBs has been suggested from amino acid sequences (Takahashi et al., 2020). First, the effect of 1 mM A22 on the swimming *Spiroplasma* cells was determined. The cells lost their original shape and stopped moving within 2 min (Fig. 2C), suggesting that the functions of SMreBs were inhibited by A22. Thereafter, we isolated the ribbon from cells maintained in 1 mM A22 for 2.5 h at 30 °C. The ribbons were found to be dispersed (Fig. 2A, c). SDS-PAGE analysis revealed contents of 67% and 11% for fibril (band iii) and SMreB2 (band v) proteins, respectively (Fig. 2B), suggesting that the protofilaments comprising fibril proteins are stabilized and modified by SMreBs in the ribbon structure.

### 3.3 Isolation and helical pitch of the fibril filament

To analyze the detailed structure of fibril filaments, we treated the ribbon fraction with cholic acid and isolated fibril proteins using sucrose-gradient centrifugation. SDS-PAGE analysis showed that the fraction only contained fibril protein (Fig. 3A). Negative-staining EM revealed that the fibril protein formed filaments that included single-, double-, and more-stranded filaments, suggesting various types of interactions between the fibril protein molecules (Fig. 3B a). A single-stranded fibril filament consisted of repeated ring units with approximately 9 nm intervals (Fig. 3B b c), consistent with previous studies (Townsend et al., 1980;Williamson et al., 1991;Trachtenberg and Gilad, 2001;Liu et al., 2017). The ring units were connected by the backbone cylinder (Fig. 3B c). The double-stranded fibril filament appeared to be formed via the alignment of two single-stranded filaments contacting with each other at the ring side not the cylinder side, resulting in a thickness of 14 nm, double that of the single-stranded filament (7 nm) (Fig. 3B d e). We analyzed the helical pitches for the double-stranded fibril filaments as the double-stranded fibril filament had a sufficient length of stable helix to cover the pitch, with a clear twist of the ring pattern along the filament axis. Images of the fibril filament cropped from the electron micrographs using the straightening selection tool of the ImageJ software were subjected to Fourier filtering to remove noise (Fig. 3C). However, the handedness of the fibril filament could not be concluded as the negative-staining EM images are projections of the object, and the alignment of the filament on the EM grid was not distinguishable. Therefore, we analyzed the isolated fibril filament using QFDE-EM (Fig. 3D) as the replica synthesized with platinum covers only one side of the object surface. The structures shared features with those from the negative-staining EM (Fig. 3D, Fig. S1). We succeeded in determining their handedness (Fig. 3D a–f) and concluded that the double-stranded fibril filament formed a left-handed helix. The half pitch was distributed at 351 ± 33 nm (n = 50), which aligns with the results of negative-staining EM (Fig. 3E). The agreement of helix pitches in the cell, isolated ribbon, and fibril filament suggests that the fibril filament is a major component of ribbon formation and cell helicity (Table 1).

**Figure 3.**
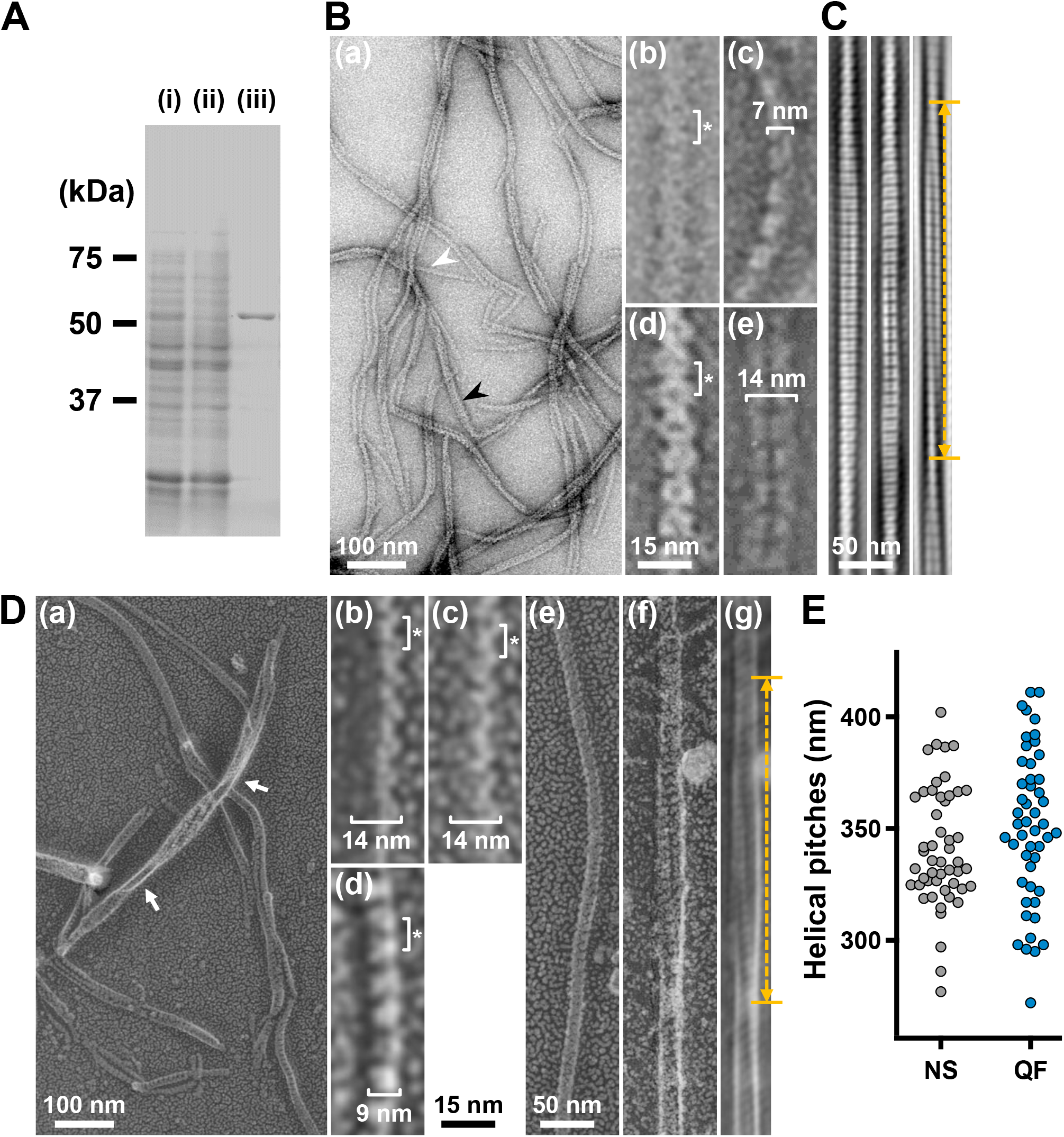
Structures of the isolated fibril filament. **(A)** Protein profiles of the fractions in the purification process for fibril protein. (i) Whole cell lysate. (ii) Supernatant. (iii) Isolated ribbon. The sample amount was adjusted to be delivered from the same cell number. **(B)** Purified fibril filaments observed by negative-staining EM. (a) Field image. White and black arrows indicate typical single and double strands, respectively. (b, c) Front and side views of the single-stranded fibril filament. (d, e) Front and side views of the double-stranded fibril filament. The ring intervals marked by an asterisk were 9 nm for both single and double strands. **(C)** Double-stranded filaments reconstituted through Fourier filtering. **(D)** Fibril filaments observed by QFDE-EM. Field (a) and single-stranded filaments (b, c, d) are presented. The back (b), front (c), and side (d) views are shown. (e) Single-stranded filament. (f, g) Double-stranded filament image (f) was reduced for noise through Fourier filtering. (g). The helical pitch was measured as depicted by a double headed broken arrow. The handedness was clearly observed at the points marked by white arrows in the panel (a) and panel (f) image.

### 3.4 Three-dimensional reconstruction of the fibril filaments

To clarify the fibril filament from a three-dimensional (3D) viewpoint, a single-particle analysis was performed on negative-staining EM images. The double-stranded fibril filament was not suitable for image averaging, which might be due to the positional variation in the binding of the two filaments (Fig. 3 and Fig. S2). Therefore, we sonicated the purified fibril fraction to increase the proportion of single-stranded filament and successfully acquired single-stranded images (Fig. 4A). From the selected 11,867 particles with good quality, the 2D-averaged images were classified into three types: (i), (ii), and (iii) (Fig. 4A b). The initial 3D image was reconstructed using the *ab-initio* 3D function of cisTEM software (Grant et al., 2018), and used as the reference for the subsequent 3D classification (Fig. 4A c). 3D structures of the fibril filament reconstructed from 11,867 particles using RELION 3.0 revealed three different conformations (i.e., class 1, left-handed mainly straight (49%); class 2, left-handed with curvature (24%); and class 3, right-handed with curvature (27%) (Fig. 4A d and Fig. S3). The class 1 structure reconstituted with rotational symmetry (C2) was not significantly different from that without symmetry (C1), suggesting that the fibril filament had rotational symmetry without polarity (Fig. S3). We therefore reconstructed the structures of the fibril filaments with C2 symmetry. The 2D reprojections from these three structures corresponded well with the 2D class averages, indicating the validity of the obtained 3D structures (Fig. S3). The 3D structure of the fibril filament had repeating elliptical rings with a pitch of 8.7 nm along the filament axis, and the ring size was 11 wide and 6 nm long along the filament. A short backbone cylinder tilted slightly to the right was found to connect the ring units, resulting in a positive curvature (Fig. 4A d). These characteristics were common to all three classes.

**Figure 4.**
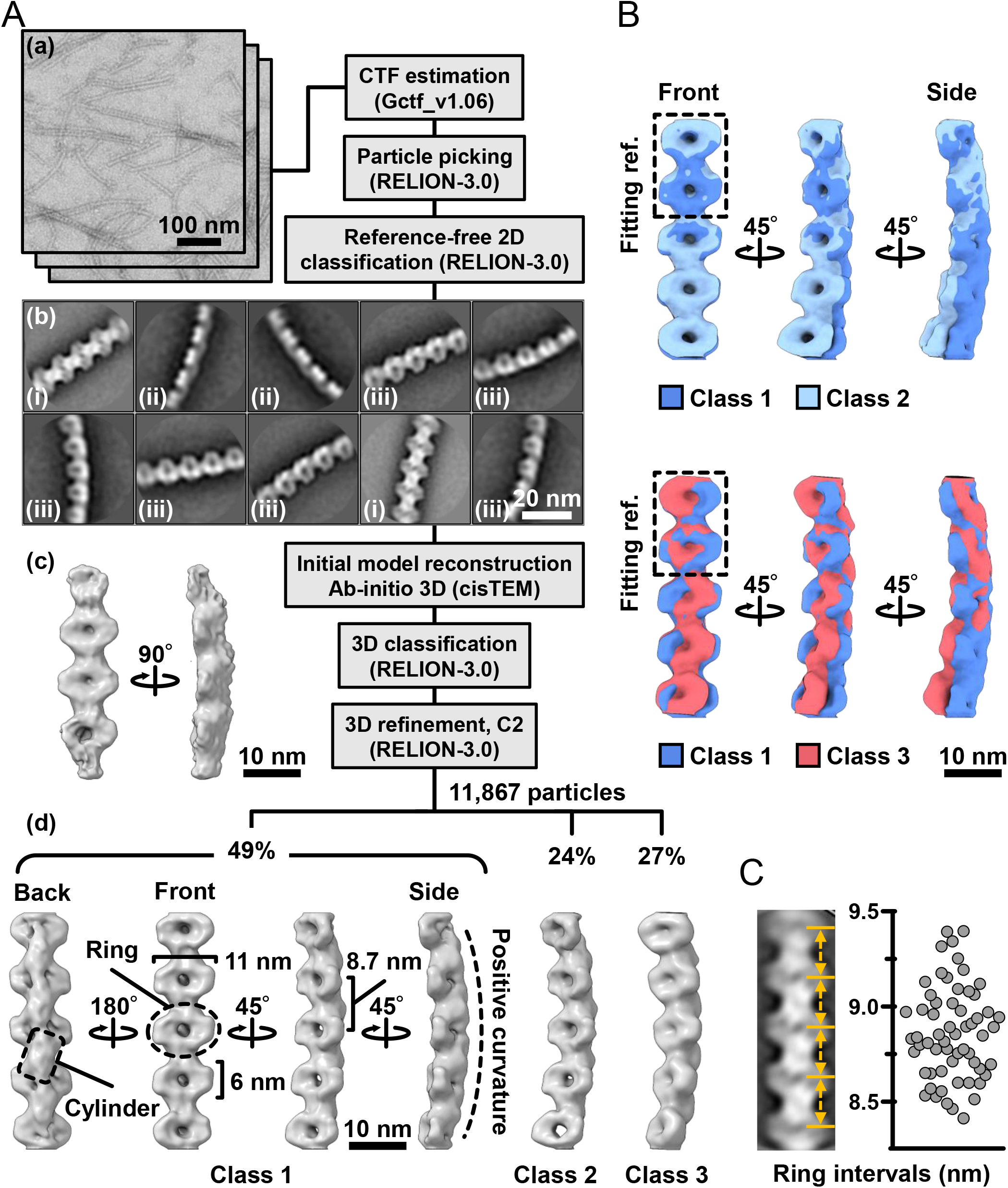
Three-dimensional reconstruction of the fibril filaments. **(A)** Workflow of single particle analysis by negative-staining EM. (a) Field images of single-stranded fibril filaments prepared by sonication. (b) Eight averaged images obtained by a function of 2D classification in RELION software. (c) The initial 3D model generated by a function of *ab-initio* reconstruction in cisTEM software. (d) Three different conformations of the fibril filament reconstituted by a function of 3D refinement in RELION software. **(B)** Superpose of class 1 (left-handed) and class 2 (left-handed) and 3 (right-handed) structures. The fitting reference is indicated by a dashed box. **(C)** Distribution of the ring intervals. Left: Ring intervals in an averaged image with complete rings. Right: Plotted ring intervals.

Although the superimposition of class 1 and others showed their structural diffferences, the positions responsible for the differences could not be identified owing to the low resolution of the structures (Fig. 4B). The fibril filaments of all classes were twisted along the filament axis, but with different rotational angles (Fig. S4). The twisting angles were estimated from the angle averages of the first and fourth units, as 5.9 (left-handed), 7.3 (left-handed), and 9.7 (right-handed) degrees for classes 1, 2, and 3, respectively. The twisting angles were estimated from the subunit numbers in the double-stranded images (Fig. 3) for negative-staining and QFDE-EM as 4.9 °and 4.7 °, respectively. These numbers slightly differed from those obtained from the reconstituted 3D structures, suggesting conformational differences between the curved and straight filament forms. These structures can explain the peak distance observed in the density profile of the isolated ribbon (Fig. 2D c, S5).

We proceeded to examine the variation in the ring interval (Fig. 4C, Fig. S6). 2D averaged images were measured for 60 ring intervals. The intervals were 8.86 ± 0.24 nm (n=60) and did not show group separation, suggesting that the intervals do not have clear conformational change, despite some having an elasticity up to 2.7%.

### 3.5 Handedness verified based on the tomography of the QFDE replica

The 3D images reconstituted from negative-staining EM had common features, despite variations in curvature and twist. The reconstructed structures all have rings and cylinders tilted slightly to the right along the filament axis when viewed from the front and back sides, respectively (Fig. 4A d), indicating that the three classes belong to the same side of mirror images. As the images by negative-staining EM are projections of the objects, the reconstituted structures may mirror images of the real structures. Thereafter, we intended to verify the handedness of the reconstituted structures by EM tomography of the QFDE replica sample (Fig. 5); this is because the tomogram cannot be a mirror image (Briegel et al., 2013;Jensen, 2015).

**Figure 5.**
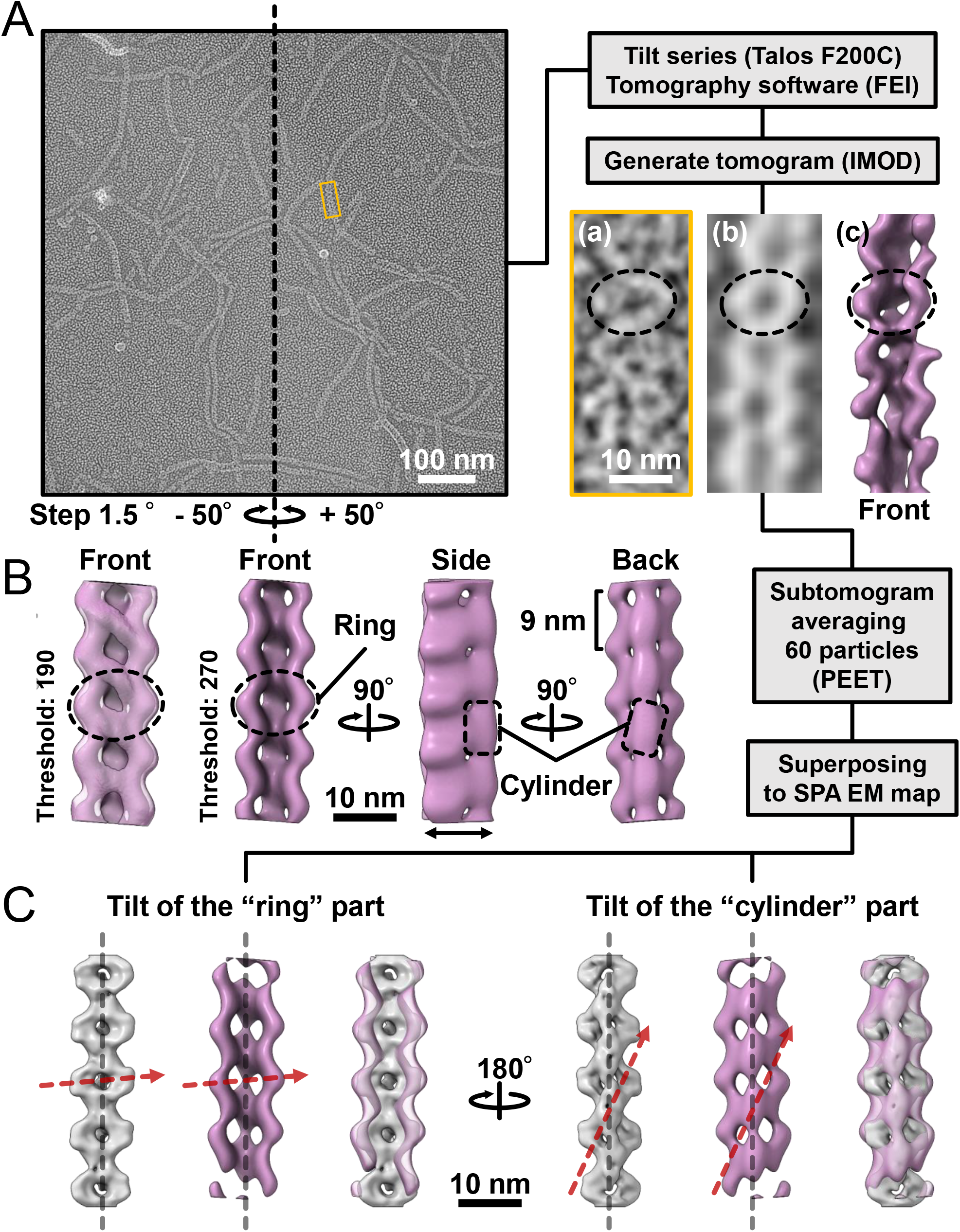
Comparison of the 3D structures of the fibril filament reconstructed from QFDE and negative-staining EM. **(A)** Replica image of mainly single-stranded fibril filaments. Left: A field image is shown from a tilt series (Movie_S2). Right lower: Magnified images of single fibril filament are shown as a raw image (a), a slice from the tomogram (b), and a subtomogram (c). **(B)** Structure averaged from 60 subtomograms. The leftmost image is presented under different thresholds from other three images. **(C)** Superpose of the 3D structures from single-particle analysis (grey) and subtomogram averaging (magenta). Long axes of ring and cylinder are depicted by broken red arrows. The filament axes were detected by a function “relion_align_symmetry --sym d2” in RELION-3.0.

We made QFDE replicas from the fraction containing single-stranded fibril filaments, acquired images every 1.5° to 50° specimen tilt for both directions, reconstituted tomograms (Movie_S4) (Fig. 5A), and then obtained a structure by averaging 60 subtomograms (Fig. 5B). As expected, the resulting filament structure had rings and cylinders. The rings and cylinders were tilted from the filament axis, rising to the right from the horizontal axis by 4–5 ° and 74–82 ° when viewed from the front and back, respectively (Fig. 5C and S7), which align well with the features of structures from negative-staining EM. These results indicate that the classes of structures from negative-staining EM had the same handedness as the real structures (Fig. 5C).

## 4 Discussion

### 4.1 Structures of the isolated fibril

The unique swimming of *Spiroplasma* is believed to be caused by the ribbon structure (Kürner et al., 2005;Cohen-Krausz et al., 2011;Harne et al., 2020b;Sasajima and Miyata, 2021). In this study, we isolated filaments of fibrils, the major protein of the ribbon, and revealed the 3D structure of the single-stranded filament at the nanometer scale using EM. Fibril filaments have been isolated for a long time, and their EM images show a characteristic ring repeat structure with a high contrast (Townsend et al., 1980;Williamson et al., 1991;Trachtenberg and Gilad, 2001;Trachtenberg et al., 2003a;Cohen-Krausz et al., 2011;Liu et al., 2017). However, a nanometer-order three-dimensional fibril filament structure is yet to be revealed. Sonication during the isolation process was effective in isolating the single-stranded filament, whose uniform structure was advantageous for image averaging (Fig. 3). Negative-staining EM was used to reconstruct the structure (Fig. 4). However, as this method produces projection images, the handedness of the reconstructed structure may be incorrect. Therefore, we confirmed the handedness of the structure by tomographic analysis of platinum replicas prepared by QFDE-EM (Fig. 5). The final structure was a repeating structure of elliptical rings connected by backbone cylinders aligned off-axis with a gentle left-handed helix, which is consistent with that of previous studies. No polarity was observed in the filament structure.

These results raise questions regarding the alignment of the 512 amino acid residues of the fibril protein with the structure and the structure formed by the 1-228 amino acid residues possessing obvious sequence similarity to methylthioadenosine/S-adenosylhomocysteine (MTA/SAH) nucleosidase (Cohen-Krausz et al., 2011;Parveen and Cornell, 2011;Sasajima and Miyata, 2021). These questions will be answered via cryo-EM analysis of the single-stranded fibril filaments prepared in this study.

### 4.2 Ribbon structure in the cell

When *Spiroplasma* cells were lysed with a detergent, the ribbon structure appeared to run along the entire length of the cell axis (Fig. 1) (Trachtenberg and Gilad, 2001). In this study, we isolated ribbons with a length equivalent with the entire length of the cell (Fig. 2). These observations suggest that the ribbon is a relatively stable structure rather than a highly dynamic one that disappears in a short time. Furthermore, as the extraction procedure with cholic acid yielded a structure consisting only of fibril filaments (Fig. 3), the stable properties of the ribbon are likely to be derived from the fibril filament. The helix of the fibril filament was directly observed in the double-stranded filament (Fig. 3). The constant helical pitch of a single strand could not be detected, which may be due to its irregular attachment to the EM grid. The two strands of double-stranded filaments may stabilize the inherent helical character of the filament by combining them. The handedness and pitch observed in the duplexes were left-handed and 351±34 (702±68) nm, respectively, aligning with the helical character of the cells at rest (Fig. 1, 3). As previous observations revealed the presence of ribbons in the innermost portion of the cell helix (Kürner et al., 2005;Trachtenberg et al., 2008), the helix of the resting cell should directly reflect the characteristics of the fibril filament.

During swimming, the cell switches its helical form into a right-handed one with a helical pitch similar to the left-handed one (Fig. 1). However, we could not find the corresponding right-handed helical structures in the isolated fibril filaments or ribbons. Only class 3 3D image reconstructed from 27% of the negative-staining EM images suggested a right-handed helical structure (Fig. 4); however, a further investigation is needed to conclude that this structure is stable one as the protein can be distorted by sticking to the EM grid in this analysis. These observations suggest that the right-handed helical structure observed in cells during swimming does not originate from another stable fibril filament structure (Fig. 4). The helix switch can also be explained by assuming that two types of filaments running parallel to the ribbon are alternately extended and contracted (Kürner et al., 2005;Cohen-Krausz et al., 2011). To test this notion, we examined the distribution of the fibril filament lengths and found that the length distribution had a single peak at 8.86 ± 0.24 nm (Fig. 4). Such finding suggests that the fibril filament has only one stable length and does not support a helical switch caused by a length change in the fibril filament.

### 4.3 Role of fibril in the swimming mechanism

The fibril protein is conserved in most *Spiroplasma* species with high amino acid sequence similarity (Ku et al., 2014). However, *Spiroplasma sabaudiense* and *Spiroplasma helicoide* do not contain fibril proteins, despite exhibiting helicity-switching swimming (Harne et al., 2020b). Recently, the expression of two SMreB proteins in the non-swimming synthetic bacterium, syn3.0B, was demonstrated to reproduce cell helicity and helicity-switching swimming (Hutchison et al., 2016; Kiyama et al., 2021). Moreover, the expression of SMreB induced cell helicity and its switching in spherical Mollicutes species (Lartigue et al., 2021), implying that the helix formation of the cell and the force generation for switching are caused by SMreBs. Then, what is the role of fibril filaments in most *Spiroplasma* species? Isolated MreB binds to fibril filaments (Harne et al., 2020a). Further, our results (Fig. 2) support the binding of SMreB to the fibril filaments. These observations suggest that MreB exerts a force on fibril filaments for swimming. SMreBs might cause helicity-switching swimming, and fibril filaments might be effective at obtaining high energy efficiency and chemotaxis; this is supported by the observation that swimming reconstructed in syn3.0B by SMreBs lacks processivity (Kiyama et al., 2021).

## 5 Conflict of Interest

The authors declare that the research was conducted in the absence of any commercial or financial relationships that could be construed as a potential conflict of interest.

## 6 Author Contributions

YS and MM designed the experiments. YS performed the experiments. YS, TK, TM, AK, and KN acquired and analyzed the images. YS and MM wrote the manuscript. All authors discussed the data.

## 7 Funding

This study was supported by Grants-in-Aid for Scientific Research A (MEXT KAKENHI, Grant Number JP17H01544), JST CREST (Grant Number JPMJCR19S5), the Osaka City University (OCU) Strategic Research Grant 2017 for top priority research to MM, JSPS KAKENHI (Grant Number JP25000013), the Platform Project for Supporting Drug Discovery and Life Science Research (BINDS) from AMED (Grant Number JP19am0101117 and support number 1282), the Cyclic Innovation for Clinical Empowerment (CiCLE) from AMED (Grant Number JP17pc0101020), and JEOL YOKOGUSHI Research Alliance Laboratories of Osaka University to KN.

## 8 Acknowledgments

We thank Yuhei O Tahara, Daichi Takahashi, Hana Kiyama, and Ikuko Fujiwara at the Graduate School of Science, Osaka Metropolitan University, Japan, for their helpful discussions.

## 1 Data Availability Statement

The datasets presented in this study can be found in the Supplementary Material.

## Notes

### Competing Interest Statement

The authors have declared no competing interest.

### Summary of Updates

Update from other studies.

