## Supplemental materials for "Isolation and structure of the fibril protein, a major component of the internal ribbon for *Spiroplasma* swimming"

Makoto Miyata

**This PDF file includes:**

Figures S1 to S7

Table S1

Legends for Movies S1 and 2

SM References

**Other supplementary materials for this manuscript include the following:**

Movies S1 to S2

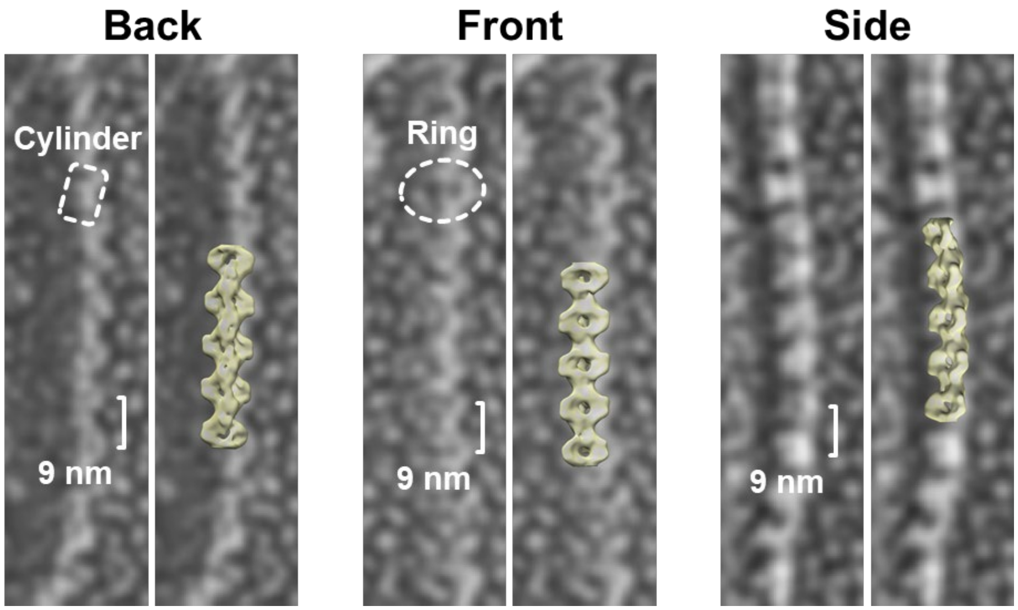


**Figure S1.** Three alignments of the double-stranded fibril filaments were observed using QFDE-EM. The features and ring intervals of 9 nm are shown in the left image of each panel. Class 1 3D images reconstituted from negative-staining EM were overlaid on the right images.

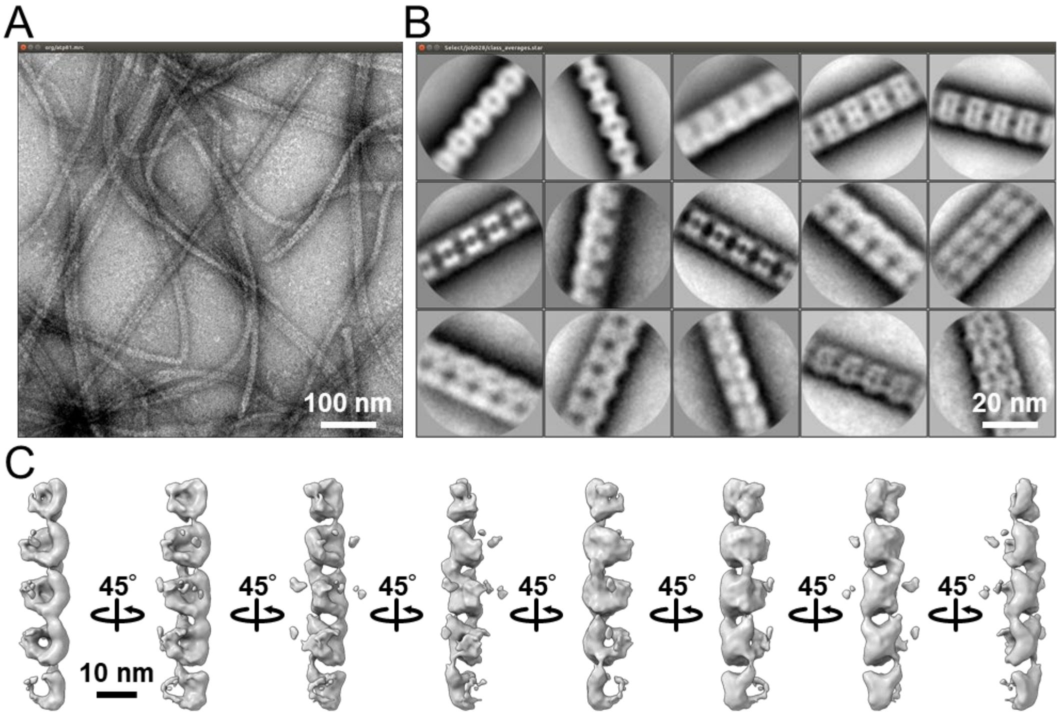


**Figure S2.** Single particle analysis using double-stranded images. (A) Field image of the isolated fibril filament observed by negative-staining EM. Most filaments displayed the double-stranded state. (B) Averaged images from the selected 16,520 segmented images constructed using "2D classification" of RELION-3.0 (1). Images of the fibril filament were segmented at a pixel size of 160 × 160, with 90% overlap as a helical object. The 15 classes of the averaged image showed different interactions between two filaments. (C) Three-dimensional reconstruction of the double-stranded fibril filament using "*ab-initio* 3D" of cisTEM (2). We failed to converge the images to solid structures.

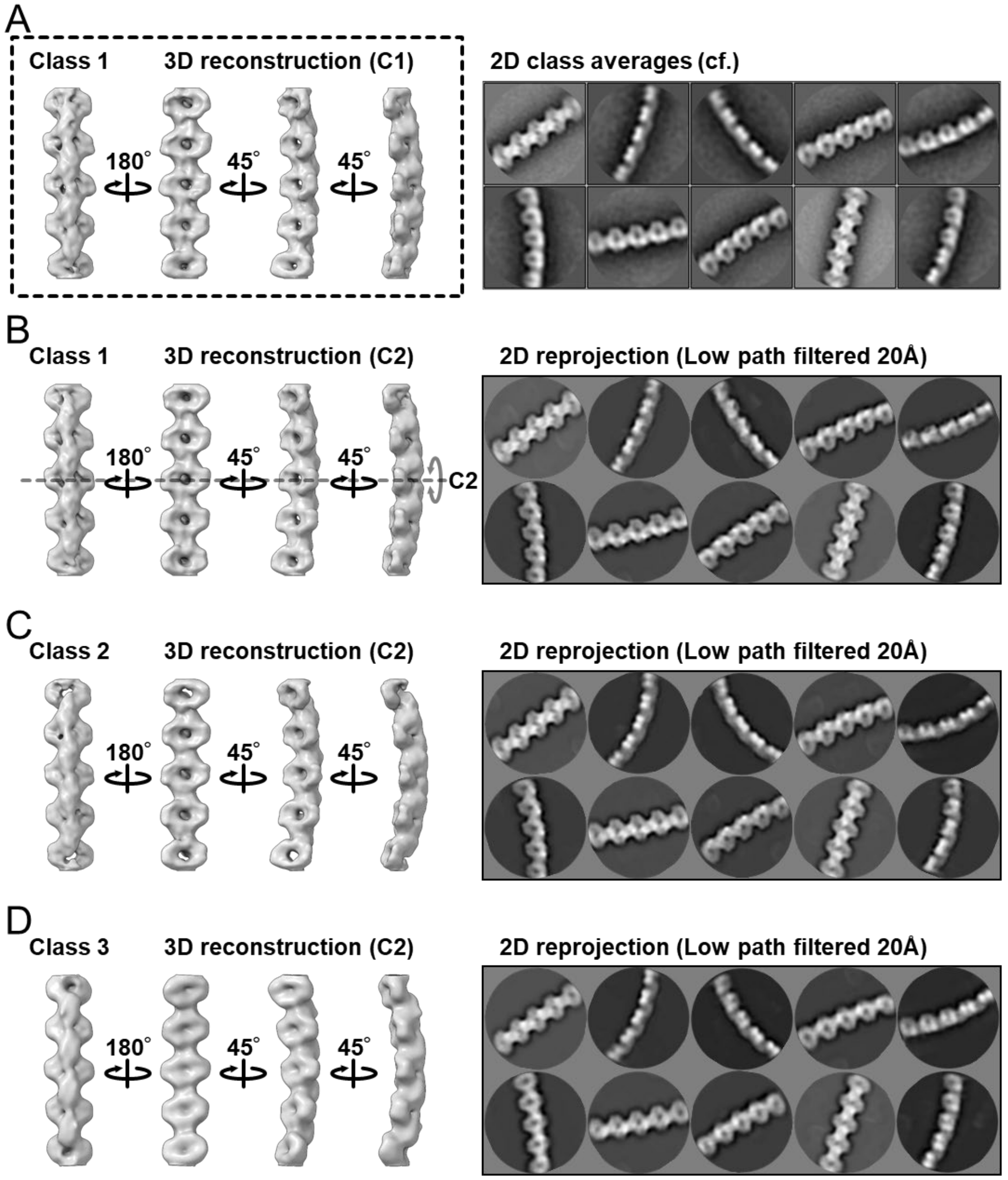


**Figure S3.** Validation of the 3D structure of the fibril filament. (A) 3D structure reconstituted as C1 from 11,867 images obtained by negative-staining EM. Individual images represent the filament views from different angles. (B-D) Three classes of 3D structures reconstructed as C2. The density map (left in each panel) and its reprojection (right in each panel) are shown as viewed at different angles. As the C2 structure of class 1 aligned well with the C1 structure, we reconstituted the structures as C2. The reprojection was processed by a lowpass filter with a resolution of 20 Å. The reprojection images appeared similar to the averaged images classified into class 1, shown at the top right, suggesting that the density maps were successfully reconstituted.

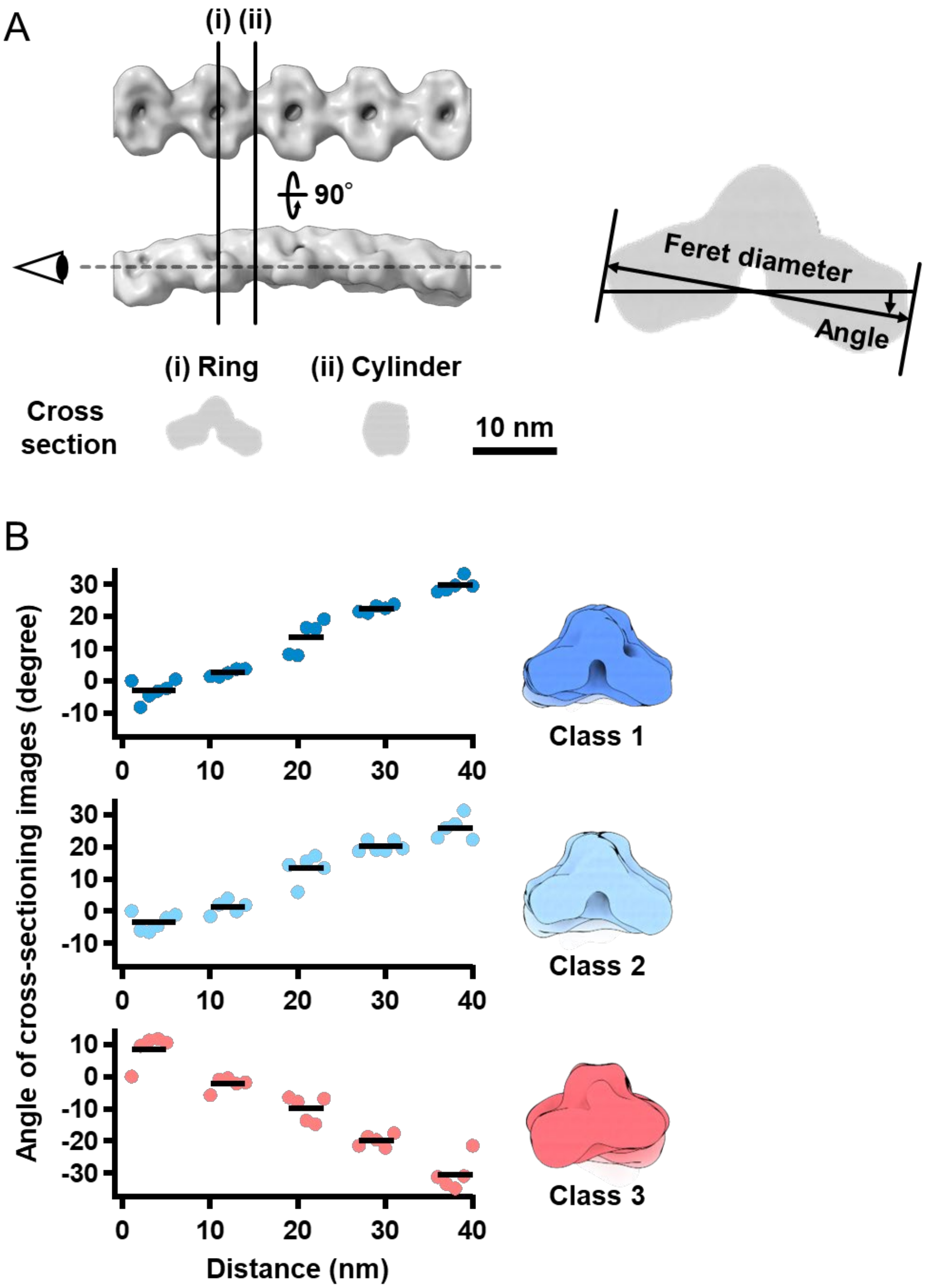


**Figure S4.** Rotation of the repeated units along the filament axis in 3D structures reconstituted from negative-staining EM images. (A) Definition of the Feret angle. The cross-sections were reconstituted from the structures for ring (i) and cylinder (ii). The alignment of the longest axis was used as the Feret angle of ring. (B) Feret angles relative to the first angle plotted along the filament. The black bars indicate positions with a Feret diameter longer than 80% of the filament maximum.

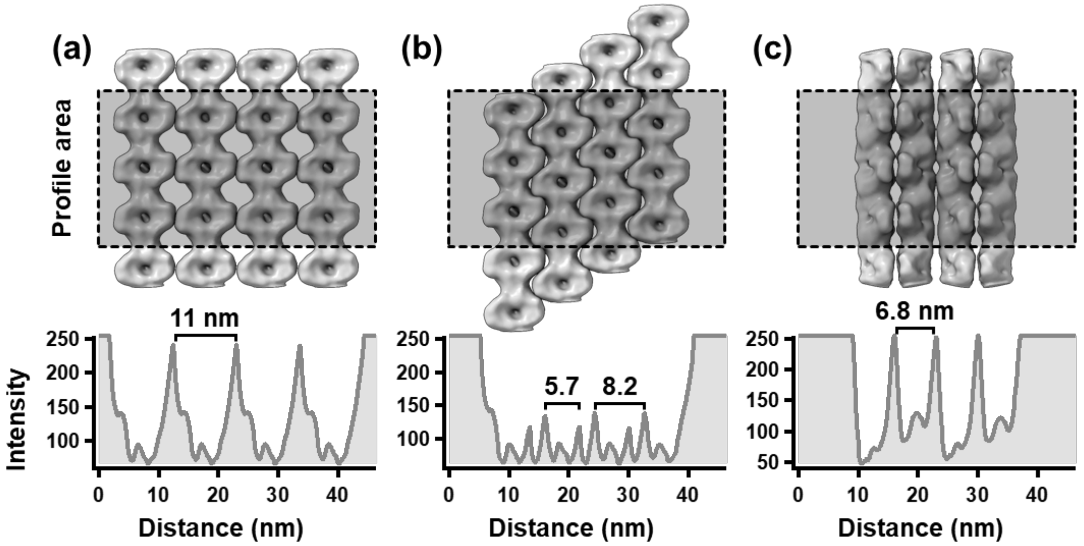


**Figure S5.** Image profile of the ribbon predicted from 3D reconstructed models of the fibril filament. Upper panels show three types of interactions between fibril filaments. Lower panels show the image profiles of the gray boxed area in the upper panels. The peak intervals are 11.0, 5.7 and 8.2, and 6.8 nm for (a), (b), and (c), respectively. The experimental values were consistent with the values of 11.0 and 6.8 nm as shown in Fig. 2D (c).

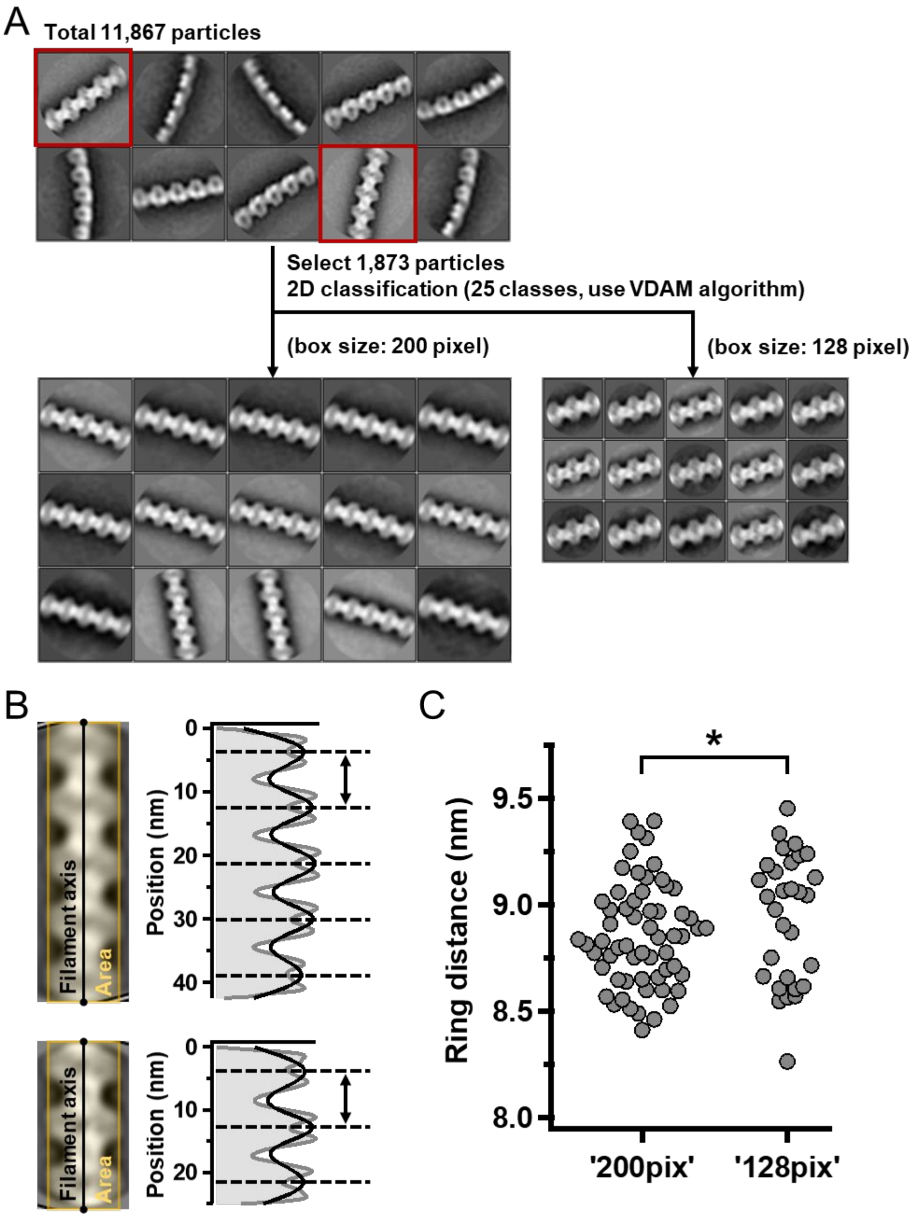


**Figure S6.** Distribution of the ring intervals. (A) 2D classification of the fibril images with complete ring structures, as marked by red boxes in ten typical images (upper). Fifteen averaged images with complete ring were selected. The selected images were derived from 1,873 images, which were used for 3D reconstruction. Two box sizes, 200 and 128 pixels, were used to estimate the effect of box size on the calculation. (B) Measurement of the ring interval. Image profile was traced for the yellow boxed area 11 nm wide along the filament axis, as shown in the left images. The profile (gray line) and gaussian fitting (black line) are shown in the right graph. (C) Distribution of the ring intervals. The results did not differ between the 200 and 128 pixels box sizes. $^{*}p > 0.05$ (the agreements of the ring intervals between the box sizes were supported by Student's *t*-test). The results obtained from 200 pixels are summarized in Fig. 4C.

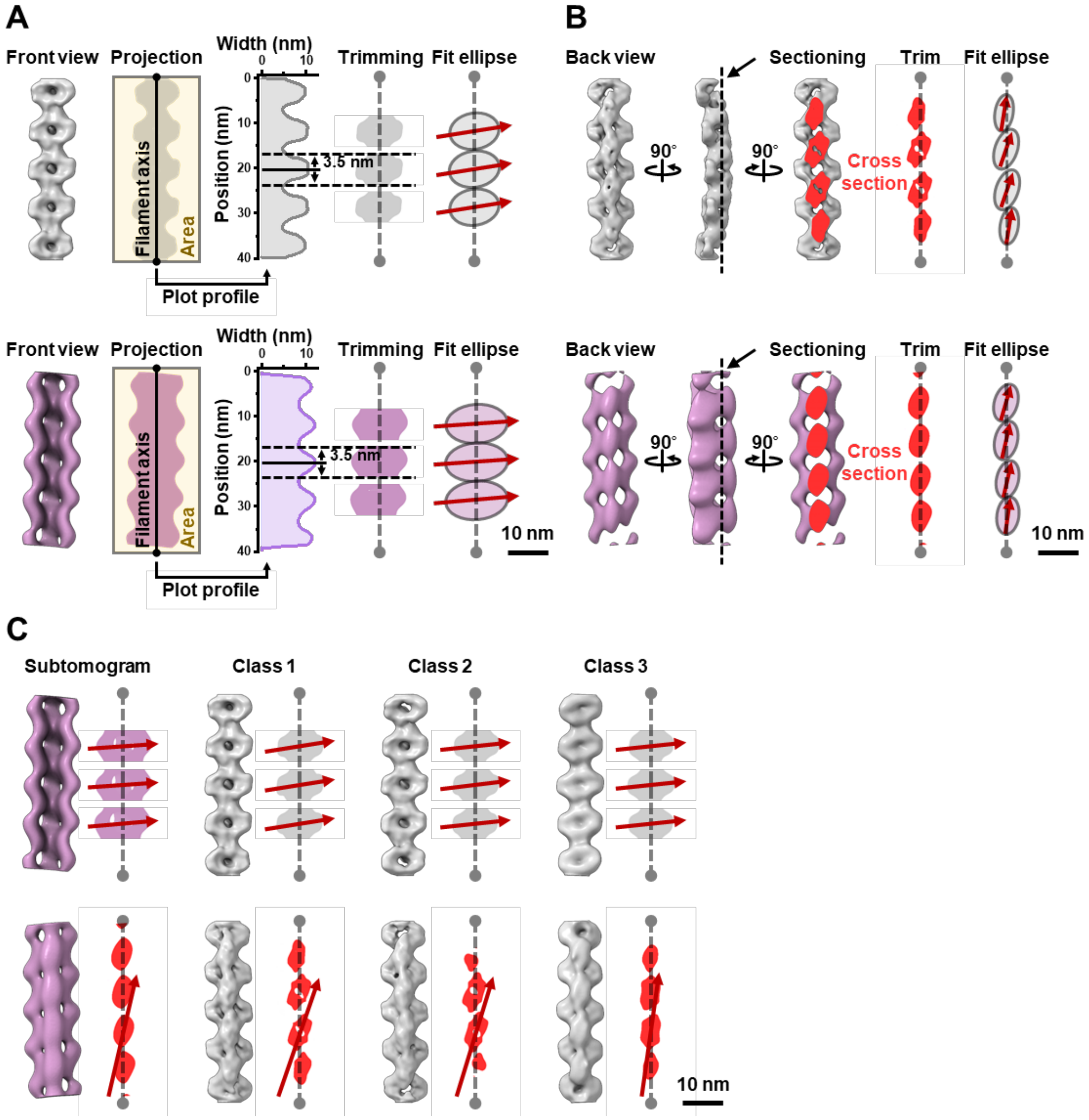


**Figure S7.** Comparison of the handedness between images obtained from negative-staining and QFDE EM. (A) Alignments of ring parts relative to the filament axis. The 3D structures reconstituted from negative-staining (class1, grey) and QFDE (magenta) EM were projected, and the width was fitted by Gaussian function to detect the peak positions. The filament axis was determined using a function “relion_align_symmetry --sym d2” in RELION-3.0. The projection image was excised as images 3.5 nm high to both sides from the peaks, and fitted as ellipses. The long axes of ellipses are indicated by red arrows. (B) Alignments of the cylinder parts relative to the filament axis. The 3D structures reconstructed from negative-staining (grey, class 1) and QFDE EM (magenta) were sectioned at the plane shown by a broken line, and the sections were fitted as an ellipse. The long axes of ellipses are indicated by red arrows. (C) Summary of the ring and cylinder alignments in reconstructed filaments from negative-staining and QFDE EM methods.

**Table S1. Peptide mass fingerprinting of band (v) in Fig. 2B.**

**SPE_1224 (MreB2)**
**Protein sequence coverage: 27%**
matched peptides are shown in ***bold red***.

1 MANYKFGKEYSFLALDLGTANTVAYVAGQGIVYNEPSMMAYDTLSNSLVA
51 LGEEAYKMIGKTHDHIKMVTPLVDGVISDMDAAQDLLK**HIFGRLK**MTGIW
101 K**NSLVILACPSGVTELER**SALKAIAKDMGASYVLVEEEVKLAALGAGINI
151 GLAQGNLVIDIGGGTTDIAILSAGDIVKSKSVKVAGKHFDQEIQKYIR**AE**
201 **YNVLIGIR**TAEQIKKDIGALVKIVNEKPIRAFGR**DIITGLPR**EVMIKPEE
251 IK**NVLLAPFSRITDLLVEVLEETPPELAGDVIRNGITICGGGALIR**GIVK
301 **YFESIFQLK**VRAAQDPLMCVIDGAKTYEKNLGAVIERIELLEAKEYKI

| Start | – | End | Mr(calc) | ppm | Peptide |
|---|---|---|---|---|---|
| **89** | **–** | **95** | **869.5235** | **-18.6** | **K.HIFGRLK.M** |
| **102** | **–** | **118** | **1856.9666** | **-31.8** | **K.NSLVILACPSGVTELER.S** |
| **199** | **–** | **208** | **1146.6397** | **-29.3** | **R.AEYNVLIGIR.T** |
| **235** | **–** | **242** | **883.5127** | **-23.2** | **R.DIITGLPR.E** |
| **253** | **–** | **261** | **1015.5815** | **-37.3** | **K.NVLLAPFSR.I** |
| **262** | **–** | **283** | **2420.3050** | **-29.9** | **R.ITDLLVEVLEETPPELAGDVIR.N** |
| **284** | **–** | **296** | **1300.6922** | **-36.8** | **R.NGITICGGGALIR.G** |
| **301** | **–** | **309** | **1173.6070** | **-32.3** | **K.YFESIFQLK.V** |

**SPE_1230 (MreB4)**
**Protein sequence coverage: 21%**
matched peptides are shown in ***bold red***.

1 **MLDIVYVYTHWK**RKKGGIFTMAGFNSGKSKRPTFVSMDLGTANTLVYVSG
51 SGVVYNEPSIVAYRIKENR**IIAVGIEAYK**MIGKGNKSIR**IVRPMVDGVIT**
101 **DIR**ATEAQLRYIFGKLRISKQLKHSIMLLACPSVITELEKAALKKIAMNL
151 GATKVFVEEEVKMAALGGGVDIYKPTGNLVVDMGGGTTDIAVIASGDIVL
201 SKSVKVAGNYLNDEMQKFIRSQYGLEVGSKTAEQIKIEIGSLAKYPDERK
251 MKVYGR**DVVSGLPREIEVTPEEVR**EVLKVPVSRIIDLTVQVLEETPPELA
301 GDIFKNGITIYGGGALIKGIDRYFTDTLQLPSK**VGEQPLLAVINGTKKFE**
351 **SDIYDILR**QEQMHTKELDY

| Start | – | End | Mr(calc) | ppm | Peptide |
|---|---|---|---|---|---|
| **1** | **–** | **12** | **1582.7854** | **27.6** | **-.MLDIVYVYTHWK.R + Oxidation (M)** |
| **70** | **–** | **79** | **1075.6277** | **-34.4** | **R.IIAVGIEAYK.M** |
| **90** | **–** | **103** | **1598.8814** | **-33.6** | **R.IVRPMVDGVITDIR.A + Oxidation (M)** |
| **257** | **–** | **264** | **841.4658** | **9.77** | **R.DVVSGLPR.E** |
| **265** | **–** | **274** | **1199.6034** | **-33.1** | **R.EIEVTPEEVR.E** |
| **334** | **–** | **347** | **1437.8191** | **-48.0** | **K.VGEQPLLAVINGTK.K** |
| **348** | **–** | **358** | **1397.7191** | **-29.6** | **K.KFESDIYDILR.Q** |
| **349** | **–** | **358** | **1269.6241** | **-27.8** | **K.FESDIYDILR.Q** |

**Movie S1 (separate file).**

*Spiroplasma* swimming was observed using phase-contrast optical microscopy. Real time. Area: 32 × 56 μm.

**Movie S2 (separate file).**

Tomography of the QFDE-EM replica from single-stranded fibril fractions. Single-axis tilt series were collected, covering an angular range from -50° to +50°, with steps of 1.5°.
